# Estimation of parental effects using polygenic scores

**DOI:** 10.1101/2020.08.11.247049

**Authors:** Jared Balbona, Yongkang Kim, Matthew C. Keller

**Affiliations:** Institute for Behavioral Genetics, University of Colorado at Boulder; Department of Psychology & Neuroscience, University of Colorado at Boulder

## Abstract

Offspring resemble their parents for both genetic and environmental reasons. Understanding the relative magnitude of these alternatives has long been a core interest in behavioral genetics research, but traditional designs, which compare phenotypic covariances to make inferences about unmeasured genetic and environmental factors, have struggled to disentangle them. Recently, Kong et al. (2018) showed that by correlating offspring phenotypic values with the measured polygenic score of parents’ nontransmitted alleles, one can estimate the effect of “genetic nurture”— a type of passive gene-environment covariation that arises when heritable parental traits directly influence offspring traits. Here, we instantiate this basic idea in a set of causal models that provide novel insights into the estimation of parental influences on offspring. Most importantly, we show how jointly modeling the parental polygenic scores and the offspring phenotypes can provide an unbiased estimate of the variation attributable to the environmental influence of parents on offspring, even when the polygenic score accounts for a small fraction of trait heritability. This model can be further extended to a) account for the influence of assortative mating at both equilibrium and disequilibrium (after a single generation of assortment), and b) include measured parental phenotypes, allowing for the estimation of the total variation due to additive genetic effects and their covariance with the familial environment. By utilizing path analysis techniques developed for extended twin family designs, our approach provides a general framework for modeling polygenic scores in family studies and allows for various model extensions that can be used to answer old questions about familial influences in new ways.

## Introduction

Parents share half of their (autosomal) additive genetic effects with their children, causing resemblance between parent and offspring for heritable traits. However, parents also help create and shape their offspring’s environment, which may have enduring influences for certain traits. If the parental traits that impact their children’s environment are themselves heritable, a correlation will develop between the genetic effects underlying those traits and the environmental effects provided by the parents. Educated parents, for example, provide to their offspring not only genes that predispose to higher education, but also a familial environment that is likely conducive to higher education. Thus, offspring who inherit genes that predispose to higher education are also more likely to be influenced by a familial environment that encourages education. This phenomenon is a type of *passive gene-environment (G-E) covariance* that has recently been referred to as *genetic nurture* (Kong et al., 2018). Generally, passive G-E covariance refers to the covariance between the genetic effects on a trait and the parenting environment influenced by that trait, regardless of whether the parenting environment actually influences the offspring. For example, aggressive parents may provide genes that influence aggression as well as an aggressive rearing environment to their children, leading to a passive G-E covariance even if parental aggression has no direct influence on offspring aggression (DiLalla and Gottesman, 1991). In such a case, an observation of a relationship between parental aggression and offspring aggression may be mistaken for a direct influence when the true cause of the association is actually shared genes. Here, we adopt the convention that *genetic nurture* refers to a specific type of passive G-E covariance that occurs when the environment provided by parents (parental aggression in this example) does directly influence the phenotype of their offspring; this leads to covariance between the genes affecting a trait and the rearing environment of the offspring influenced by that trait in the parents. Genetic nurture is therefore a necessary consequence of the direct effect of parental phenotype on offspring phenotype, known as *vertical transmission* (VT) in the behavioral genetics literature (Cavalli-Sforza and Feldman, 1973). While passive G-E covariance could also be due to *horizontal transmission*— in which other collateral relatives (typically siblings) influence one another— we focus in this paper on genetic nurture arising from VT and discuss horizontal transmission at the end.

The occurrence of genetic nurture has important implications for understanding complex trait genetics. Genetic nurture increases the phenotypic variation in the population over what it would otherwise be, and can bias estimates of genetic or environmental influences. In genome-wide association studies (GWAS’s), genetic nurture inflates estimates of SNP associations. This bias can be quantified in part by comparing within-family estimates, which are immune to this inflation, to standard between-family estimates. For example, Lee et al. (2018) found that SNP associations with educational attainment estimated within-families were 40% smaller than those estimated between-families, consistent with the presence of genetic nurture. Similarly, by inflating the associations between traits and their causal variants across the genome, genetic nurture upwardly biases estimates of SNP-heritability, including those from methods that use GWAS summary statistics (e.g., LD-score regression; Bulik-Sullivan et al., 2015) and from methods that use average similarity across genome-wide SNPs (e.g., genomic REML; Yang et al., 2011). Genetic nurture also increases dizygotic and monozygotic twin covariances to the same degree, leading to overestimates of the shared environmental variation in the classical twin design “ACE” models, and (less intuitively) to overestimates of the additive genetic variance in “AE” or “ADE” models that estimate only genetic variation (Coventry and Keller, 2005). For these reasons, quantifying genetic nurture is important for interpreting estimates across multiple designs and approaches.

Detecting genetic nurture using traditional family-based designs has been challenging because no observed statistic directly provides information to estimate it. Genetic nurture must rather be inferred as a consequence of estimated heritability that co-occurs with estimated VT (Eaves, 1976). In turn, VT must be estimated from residual parent-offspring covariance that is higher than expected from estimated genetic and environmental influences. However, many factors such as dominance, epistasis, and background environmental effects can influence parent-offspring covariances, and these cannot be simultaneously modeled. Most existing estimates of genetic nurture and VT thus depend strongly on model assumptions. Recently, Kong et al. (2018) demonstrated an innovative approach for the near-direct estimation of genetic nurture using measured polygenic scores (PGS’s; the predicted genetic scores for a trait based on SNP weights from a GWAS conducted in an independent sample). From data on ∼ 22K Icelandic offspring and their parents in the deCODE sample, Kong et al. constructed two groups of PGS’s for educational attainment: one using the set of alleles parents transmitted to their offspring, and one using the set of alleles parents did not transmit to their offspring. After controlling for the potential confounding influences of stratification, they found a highly significant relationship between the nontransmitted PGS and offspring educational attainment. Such an association should not occur if the genetic and environmental causes of parent-offspring resemblance are independent of one another. Rather, if the effects of potential confounding factors such as stratification and assortative mating (AM) are properly accounted for, it is difficult to imagine alternative explanations for this association beyond genetic nurture due to VT.

The significance of the Kong et al. paper is less that it introduced new concepts, but that it introduced a new and better way to test old ones. While their idea of using PGS’s to understand genetic nurture is an extremely important advance in family studies and behavioral genetics, we identify three aspects of their approach that we attempt to improve upon here. First, while genetic nurture is interesting in its own right, it is not the only construct of interest that can be estimated from parental PGS’s. As noted above, the observation of genetic nurture implies the presence of VT, which Kong et al. did not estimate. Thus, we demonstrate below how a causal model that incorporates parental PGS’s and offspring phenotypes can be used to estimate the total variation accounted for by VT (*V*_*F*_), even when the PGS accounts for a small fraction of trait heritability. Second, most of the information to estimate *V*_*F*_ and genetic nurture comes from the observed relationship between the nontransmitted PGS and the offspring phenotype. However, primary phenotypic AM (hereafter simply ‘AM’)—the tendency for parents to chose mates who are similar to themselves—is a competing explanation for this relationship that must be accounted for. Because Kong et al. found evidence that AM for educational attainment in Iceland occurred only in the parental generation and not before, their approach assumed this specific type of disequilibrium AM. Using techniques developed for extended twin family designs, we demonstrate how both disequilibrium and equilibrium AM can be tested and incorporated in causal models. Finally, much of the math presented by Kong et al. was derived from first principals. While an impressive feat, this limits the generalizability of their approach. Here, we present a simple causal modeling framework for using PGS’s in family studies that allows these models to be extended in ways that would otherwise lead to intractable math. In that vein, we extend the models to include parental phenotypes, which allow for unbiased estimates of the total variation due to additive genetic effects, genetic nurture, and *V*_*F*_, all while accounting for equilibrium and disequilibrium AM.

The set of models presented here can be used to better understand the genetic and environmental influences of parents on offspring. While they can be implemented “out of the box”, their greater utility is as examples of a general approach for incorporating PGS’s into family causal models in order to test specific hypotheses of interest. To this end, we present the logic and math underlying the models, adopting a tutorial style so that the models presented can be modified and improved upon to best fit the trait and data being worked with.

### Overview of path analysis

To aid in the understanding of our models, we first review the core concepts underlying path analysis as used to model extended twin family data in the behavioral genetics literature. Path analysis was initially developed by Sewell Wright (Wright, 1934) as a method of deriving the expected variances and covariances among variables in a particular *causal model* — the set of assumed causal connections between measured and unmeasured variables. Although sometimes misunderstood, path analysis does not test causation but rather assesses the degree to which a set of observed variances and covariances is consistent with a particular causal model (Bollen and Pearl, 2013). To this end, models are fit using structural equation modeling software, such as OpenMx (Boker et al., 2011), which provide estimates that maximize the likelihood of the data given the assumed causal model.

Path analysis is very useful in the present context for several reasons. First, it provides a set of rules that greatly simplify finding mathematical expectations of variances and covariances under potentially complicated scenarios, such as those involving VT and AM. Second, path analysis encourages a focus on effect sizes (estimated variances and covariances) and their standard errors rather than on hypothesis tests and *p*-values. Third, it uses the data efficiently, finding estimates that account for all other relationships simultaneously instead of finding estimates in only certain subsets of the data at a time. Fourth, path analysis forces the user to think carefully about the possible causal mechanisms that underlie observed data. Lastly, and related to the previous point, path analysis requires that the user make model assumptions explicit and (hopefully) encourages testing these assumptions. This final advantage is related to a weakness of path analysis: the estimates it produces depend to various degrees on the assumptions made when building the causal model. To the degree that assumptions are unmet, estimates will be biased. Therefore, to properly interpret estimates from a given model, it is crucial to understand how estimates behave under different scenarios in which some assumptions are unmet; for complex models, this typically requires comparing estimates to known parameters in simulated data, which we do for the models below in our companion paper (Kim et al., 2020, under review).

A causal model is considered ‘identified’ if there is a unique solution for all the model’s parameters. Often, there is insufficient information to estimate all parameters of interest, requiring that the values of some parameters be fixed (typically to 0 or 1) so that the model is identified; however, it should also be noted that fixing parameters is itself a model assumption. Even for models that are identified, certain assumptions about causal relationships between variables can lead to high correlations among estimates, increasing their standard errors and, in some cases, necessitating that one or more of the parameters be fixed.

Path diagrams pictorially represent causal models and are helpful for deriving the variances and covariances that the models imply. By convention, squares represent observed variables and circles represent unobserved ‘latent’ variables— theorized causes of variation and covariation among observed variables. Single-headed arrows from one variable to another signify causal relationships from the former to the latter, with their associated path coefficients being akin to partial regression coefficients. Double-headed arrows, meanwhile, signify covariance when connecting two variables to each other and variance when connecting a variable to itself. Finally, a straight line with no arrows is called a ‘copath’ and is used to model AM. The copath is a more recent innovation in path analysis (Cloninger, 1980) and is not widely used outside the extended twin family literature.

To derive expectations using a path diagram, one must identify all legitimate paths that connect two variables (for expected covariances) or that connect a variable back to itself (for expected variances). These paths can be thought of as chains, with each individual arrow or copath representing a link in a particular chain. The expected value of a chain is equal to the product of all the coefficients associated with each of its links, with the final expected variance or covariance being equal to the sum of all legitimate chains. A chain is considered legitimate if it abides by the following path tracing rules:

1. A chain begins by travelling backwards against the direction of a single or double-headed arrow (from the arrow’s head to its tail). However, once a double-headed arrow has been traversed, the direction reverses such that the chain now travels forwards, in the direction of the arrows.
2. A chain must include exactly one double-headed arrow (a variance or a covariance term), which is equivalent to stating that a chain must change directions exactly once. This is necessary because double-headed arrows provide the proper scaling for the coefficients in each chain.
3. All chains must be counted exactly once and each must be unique. However, the order of the links in the chains matters. For example, despite being algebraically equivalent, the chain *Y*_*p*_ → *NT*_*p*_ → *T*_*p*_ → *Y*_*p*_ is distinct from the chain *Y*_*p*_ → *T*_*p*_ → *NT*_*p*_ → *Y*_*p*_ in Figure 1. Both are unique and both must be counted in determining the variance of *Y*_*p*_
4. Copaths may only be traversed once in a given chain, and a chain must be legitimate before traversing the copath. However, once the copath is crossed, the first two rules above reset. A chain must therefore contain exactly one double-headed arrow before traversing the copath, and one double-headed arrow after traversing the copath. Thus, copaths connect two legitimate chains to create a single, longer chain.

To demonstrate the first three rules (the fourth is demonstrated below), we derive the expected *cov*(*Y*_*p*_, *NT*_*p*_) in Figure 1, denoted as Ω in our models. As mentioned, deriving the covariance between two terms requires tracing all legitimate chains that connect them. In this case, only two legitimate chains start at *Y*_*p*_ and end at *NT*_*p*_ (though one could equivalently start at *NT*_*p*_ and end at *Y*_*p*_). The first travels up the arrow *Y*_*p*_ → *NT*_*p*_ (path coefficient *δ*), and because all chains require a double-headed arrow, finishes by traversing the double-headed arrow leading back to *NT*_*p*_ (i.e., the variance of *NT*_*p*_, with path coefficient 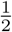). The second travels up the arrow *Y*_*p*_ → *F*_*p*_ (with path coefficient 1) and then traverses the double-headed arrow *F*_*p*_ → *NT*_*p*_ (i.e., the covariance between *F*_*p*_ and *NT*_*p*_, with path coefficient 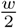). Thus, 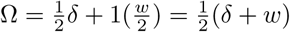.

**Figure 1.**
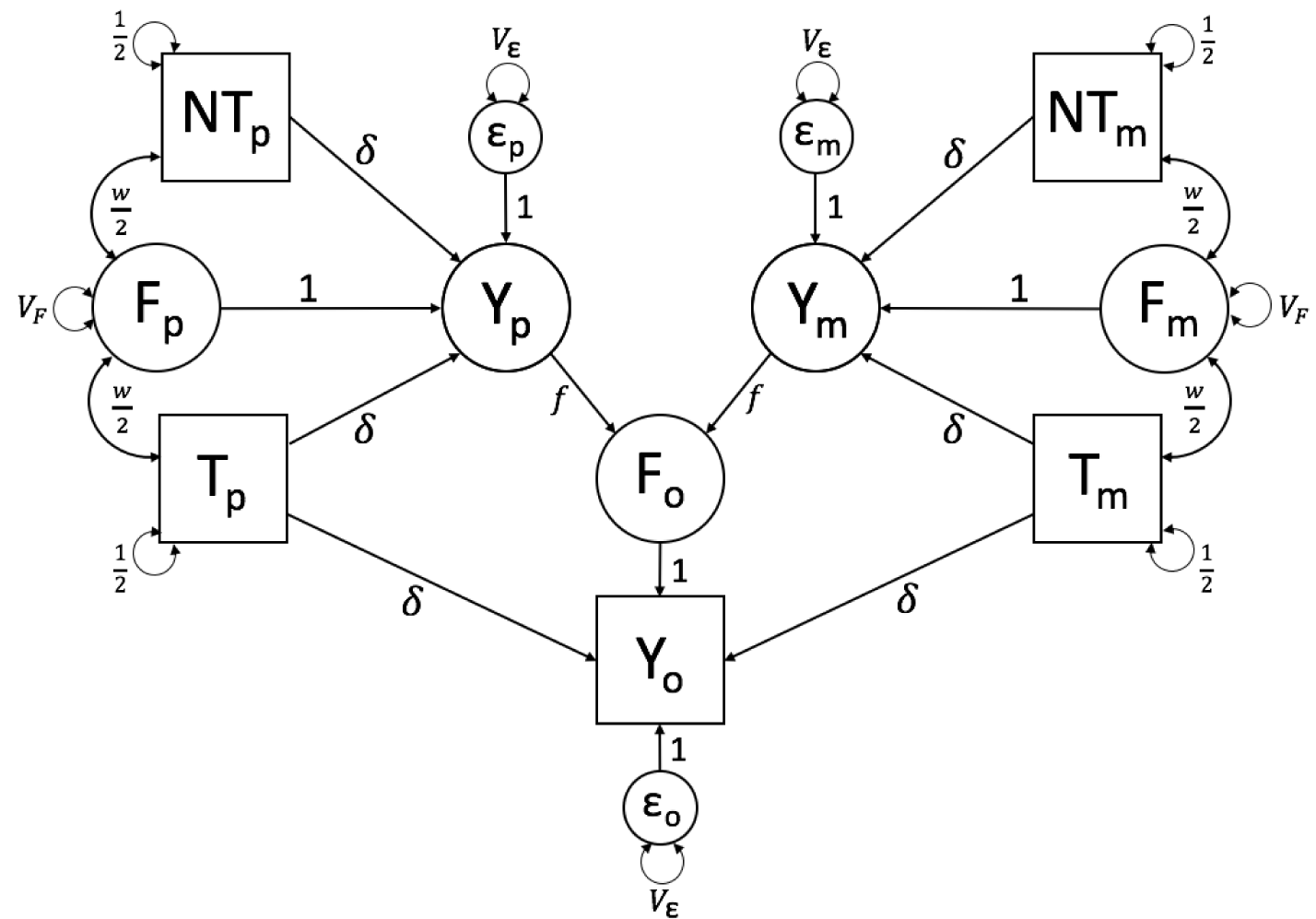
Path diagram of Model 0, which models the effects of VT, assuming that the PGS explains the full trait heritability and that there is no AM

### Overview of the models

Although the path tracing procedures described above are simple and algorithmic, the number of unique chains grows rapidly as models become more complicated, making the process error-prone. To simplify chains and reduce the probability of errors, we substitute variables (e.g., Ω) for chain segments that recurred across multiple chains 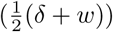 when possible. We use parameter subscripts *p, m*, and *o* for paternal, maternal, and offspring, respectively. To reduce redundancy, we use [*N*]*T* to denote either *NT* (the nontransmitted) or *T* (the transmitted) haplotypic PGS, and we use the ∗ subscript to denote either *p* or *m* but not both within a single term. For example, *cov*(*Y*_∗_, [*N*]*T*_∗_) can be written in the place of *cov*(*Y*_*p*_, *T*_*p*_), *cov*(*Y*_*p*_, *NT*_*p*_), *cov*(*Y*_*m*_, *T*_*m*_), or *cov*(*Y*_*m*_, *NT*_*m*_). However, *cov*(*Y*_∗_, *T*_∗_) does not equal *cov*(*Y*_*p*_, *T*_*m*_) or *cov*(*Y*_*m*_, *T*_*p*_), which mix *m* and *p* within the same term. Finally, to ease comparison, we follow the notation of Kong et al. when possible. For example, the meanings of *δ, θ*_*T*_, *θ*_*NT*_, *T*_*p*_, *NT*_*p*_, *T*_*m*_, and *NT*_*m*_ are consistent across papers. The names of other parameters (*V*_*F*_, *w, f, V*_*E*_, and *µ*) were chosen for consistency with existing extended twin family models. For descriptions of these and other parameters, see Table I.

**Table I:** Parameters in Models 0, 1, and 2.

| Parameter | Interpretation |
| --- | --- |
| $Y$ | phenotypic score |
| $V_Y$ | phenotypic variance |
| $\epsilon$ | residual score |
| $V_\epsilon$ | residual variance not explained by other factors |
| $\mu$ | AM (assortative mating) copath coefficient; $\frac{cov(Y_p, Y_m)}{V_Y^2}$ |
| $F$ | family environmental score arising from $Y_* \rightarrow F_o$ vertical transmission (VT) |
| $f$ | VT path coefficient; the causal effect of $Y_* \rightarrow F_o$ |
| $V_F$ | familial environmental variance caused by VT |
| $T_*$ | polygenic score (PGS) of one of the two transmitted haplotypes |
| $NT_*$ | PGS of one of the two nontransmitted haplotypes |
| $k$ | variance of the haplotypic PGS in the base population (before AM or VT). It is a constant that depends on the scaling of the PGS (see Model 1) |
| $g$ | increase in the (co)variance of the haplotypic PGS's under AM |
| $\delta$ | effect of haplotypic PGS on $Y$ |
| $w$ | genetic nurture between the PGS and $F$; $cov(T_* + NT_*, F_*) = cov([N]T_p + [N]T_m, F_o)$ |
| $\Omega$ | covariance between $Y_*$ and either of that parent's haplotypic PGS's; $cov(Y_*, [N]T_*)$ |
| $\theta_T$ | covariance between $Y_o$ and both of the transmitted haplotypes; $cov(Y_o, T_m + T_p)$ |
| $\theta_{NT}$ | covariance between $Y_o$ and both of the nontransmitted haplotypes; $cov(Y_o, NT_m + NT_p)$ |
| $LT_*$ | latent genetic score (LGS) of one of the two transmitted haplotypes |
| $LNT_*$ | LGS of one of the two nontransmitted haplotypes |
| $j$ | variance of the haplotypic LGS in the base population. It is a constant defined analogously to $k$ (see Model 2) |
| $h$ | increase in the (co)variance of the haplotypic LGS's under AM |
| $i$ | increase in the covariance between the haplotypic PGS's and LGS's under AM |
| $a$ | effect of haplotypic LGS on $Y$ |
| $v$ | genetic nurture between the LGS and $F$; $cov(LT_* + LNT_*, F_*) = cov(L[N]T_p + L[N]T_m, F_o)$ |
| $\Gamma$ | covariance between $Y_*$ and either of that parent's haplotypic LGS's; $cov(Y_*, L[N]T_*)$ |
| $\theta_{LT}$ | covariance between $Y_o$ and the transmitted LGS; $cov(Y_o, LT_m + LT_p)$ |
| $\theta_{LNT}$ | covariance between $Y_o$ and the nontransmitted LGS; $cov(Y_o, LNT_m + LNT_p)$ |

Subscripts:
$m$ : maternal; $p$ : paternal; $o$ : offspring; $*$ : $m$ or $p$ ; no subscript: $m$ , $p$ , or $o$

To ensure that our parameter derivations are correct, we compared expected equilibrium parameter values to simulated ones from an adapted version of the *GeneEvolve* software (Tahmasbi and Keller, 2017), which we modified for efficiency such that causal variants were in linkage equilibrium in the *base population*—the population before any AM or VT has occurred. Because most parameters depend on each other recursively, we found their expected equilibrium values by inputting start values into their expectations derived below and iterated all parameters together in R until their values converged. For all models, the expected equilibrium values of the parameters agreed with their observed equilibrium counterparts from *GeneEvolve*.

Each of our models require four pieces of information: the PGS of the mother, the PGS of the father, the offspring phenotypic value, and the offspring genotype (in order to differentiate the transmitted and nontransmitted parental alleles). We show that this information is sufficient for estimating the full *V*_*F*_ when there is no AM (Model 0). However, AM induces covariances between latent and observed genetic scores, biasing estimates of *V*_*F*_ and genetic nurture when the PGS accounts for little of the heritability (Model 1). This bias can be eliminated by modeling the genetic variation not captured by the PGS (Model 2), either through estimating it by including parental phenotypes in the model or by making an assumption about its value in the base population. Of course, each of these models require assumptions; we describe these in each subsection, but focus on their influences in a companion paper (Kim et al., 2020, under review).

### Model 0: VT but no AM

Figure 1 shows a path diagram of the simplest model of genetic nurture and so serves as a valuable starting place. It makes two assumptions that distinguish it from later models: 1) there is no AM, and 2) the PGS explains all the genetic variation in the trait. The first assumption will be unmet for many traits of interest while the latter assumption is unmet for all traits currently. Nevertheless, when the first assumption is met (no AM), we show below that this simple model can provide unbiased estimates of the full *V*_*F*_.

This model estimates five unknown parameters: *δ*, the direct effect of haplotypic PGS on the phenotype after removing the influence of genetic nurture; *f*, the direct effect of parental phenotype on the offspring environment (i.e., the VT effect) ; *V*_*F*_, the familial variance due to VT; *w* the genetic nurture effect; and *V*_*E*_, the variance of the residual phenotypic variation. It is worth noting that the values of *f* and *V*_*F*_ are determined given the values of *δ, w*, and *V*_*ϵ*_, and so only three of these five estimates are independent. Additionally, the parental phenotypes (*Y*_*p*_ and *Y*_*m*_), familial environment value arising from VT (*F*), and unique environmental score (*ϵ*) are latent and are therefore represented by circles. To prevent under-identification, the *F* → *Y* and *E* → *Y* paths are fixed to 1. Similarly, the variances of the haplotypic PGS’s are constrained to 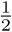, which should be true if the full PGS is standardized and there is no AM to induce covariances between haplotypic PGS’s.

As previously stated, there are five observed variables in this model—the transmitted and nontransmitted paternal (*T*_*p*_ and *NT*_*p*_) and maternal (*T*_*m*_ and *NT*_*m*_) haplotypic PGS’s as well as the offspring phenotype (*Y*_*o*_)—creating a 5-by-5 observed variance-covariance matrix and leading to 15 unique statistics from which to estimate parameters. Model-fitting software mimics as closely as possible this observed variance-covariance matrix with the one implied by the maximum likelihood estimates of the unknown parameters. While 15 independent statistics is easily sufficient for estimating a model with three unknowns, many of the statistics in this model provide redundant information. The four haplotypic PGS variances and the six covariances between them are assumed to be constants (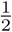 and 0, respectively) and provide no information for estimating parameters. The remaining five statistics provide only three independent pieces of information: one from the two covariances between the haplotypic nontransmitted PGS (*NT*_∗_) and *Y*_*o*_, one from the two covariances between the haplotypic transmitted PGS (*T*_∗_) and *Y*_*o*_, and one from the variance of *Y*_*o*_. These three independent sources of information are used to estimate three independent parameters (*δ, w*, and *V*_*ϵ*_). Thus, this model is just-identified.

Although parental phenotypes are unobserved in this model, it is still useful to define the covariance between haplotypic PGS’s and the latent parental phenotypes because this term recurs throughout. We denote this covariance as Ω and, as noted above, 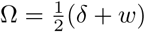. Under this model’s assumptions of no sex-specific genetic or VT effects, Ω is the same regardless of the PGS’s parental origin or whether it is transmitted: *cov*(*T*_*p*_, *Y*_*p*_) = *cov*(*NT*_*p*_, *Y*_*p*_) = *cov*(*T*_*m*_, *Y*_*m*_) = *cov*(*NT*_*m*_, *Y*_*m*_). Thus, Ω can be used as a substitute for 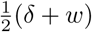 in any chain that traverses *Y*_∗_ → [*N*]*T*_∗_ or [*N*]*T*_∗_ → *Y*_∗_ in order to simplify finding other expected values, such as the two covariances at the core of this model:

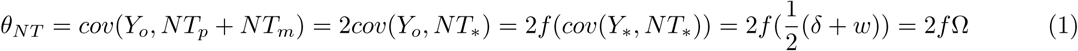

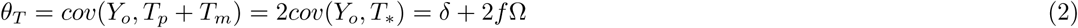

Kong et al. emphasized that part of the relationship between *Y* and its PGS (*T*_*p*_ + *T*_*m*_) may be due to the confounding influences of genetic nurture. This can be seen in the additional 2*f* Ω term in *θ*_*T*_ above. Thus, as noted by Kong et al., *θ*_*T*_ − *θ*_*NT*_ = *δ* is an estimate of the direct genetic effect of the PGS, controlling for genetic nurture.

This model assumes that parameters have reached equilibrium, which implies that variances and covariances are the same across parental and offspring generations. The equilibrium assumption allows the parameters that change over time (*V*_*Y*_, *w*, and *V*_*F*_) to be estimated by constraining their values in the parental generation to their derived values in the offspring generation. For example, at equilibrium, the covariance between *F*_∗_ and the haplotypic PGS’s in the parental generation 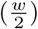 must equal the implied covariance between *F*_*o*_ and any of the four haplotypic PGS’s (*f* Ω, which can be found through path tracing). Thus,

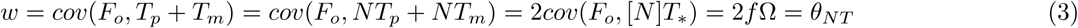

Note that this estimated value of *w* is equal to the estimated value of *θ*_*NT*_ derived in equation (1), indicating that *θ*_*NT*_ is a direct estimate of genetic nurture (under the assumption of no AM). Meanwhile, the variance of *Y*_*p*_ (denoted by *V*_*Y*_) is derived by summing all chains that begin at *Y*_*p*_ and end back at *Y*_*p*_, and is assumed to be equal to the variance of *Y*_*m*_ and *Y*_*o*_:

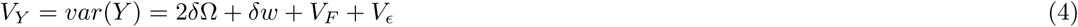

Finally, the expectations for the variances of *F*_*p*_ and *F*_*m*_ (*V*_*F*_) can be found by constraining their values to all legitimate chains that connect *F*_*o*_ back to itself, of which there are two: (1) *F*_*o*_ → *Y*_*p*_ → *Y*_*p*_ → *F*_*o*_ and (2) *F*_*o*_ → *Y*_*m*_ → *Y*_*m*_ → *F*_*o*_. Thus,

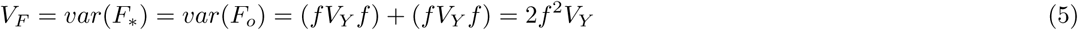

Note that the variance *F*_*o*_— as well as its covariances with the haplotypic PGS’s— are not shown in any of the models, as it is already implied through the connections between *F*_*o*_ and the parental phenotypes; explicitly including *V*_*F*_ and 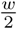 in the offspring generation would thus be redundant, resulting in expectations that are double their true values. Furthermore, note that *V*_*F*_ is a function of *V*_*Y*_ but that *V*_*Y*_ is also a function of *V*_*F*_. Similarly, Ω is a function of *w*, which is a function of Ω. These types of recursive relationships are known as *nonlinear constraints*, which describe and constrain such interdependent relationships between parameters in a way that keeps the overall model internally consistent and identified. The implementation of nonlinear constraints is a hallmark of family models, and requires the use of optimizers (such as NPSOL in OpenMx) that can estimate their values iteratively.

Last, we show that when the assumption of no AM is met, this model provides a full estimate of *V*_*F*_, regardless of the amount of variance explained by the PGS (*δ*^2^). Given equation (1), as well as the knowledge that *w* = *θ*_*NT*_ and *δ* = *θ*_*T*_ − *θ*_*NT*_,

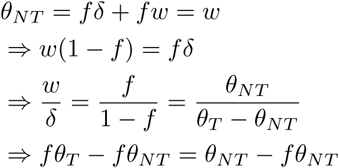

Through rearrangement of terms,

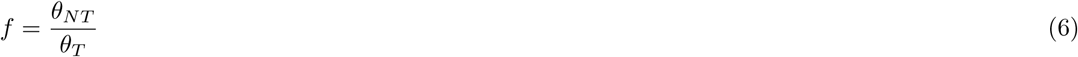

Thus, the estimate of *V*_*F*_ (= 2*f* ^2^*V*_*Y*_) depends on only three observed statistics: *θ*_*NT*_, *θ*_*T*_, and *V*_*Y*_. Note that the expectation of *θ*_*NT*_ contains two parameters, *w* and Ω, that are functions of one another. Substituting the value of *w* recursively thus leads to a geometric series:

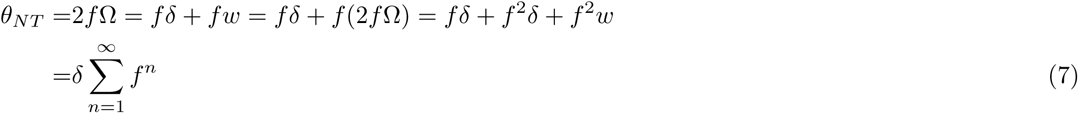

Therefore, because *θ*_*T*_ = *δ* + *θ*_*NT*_,

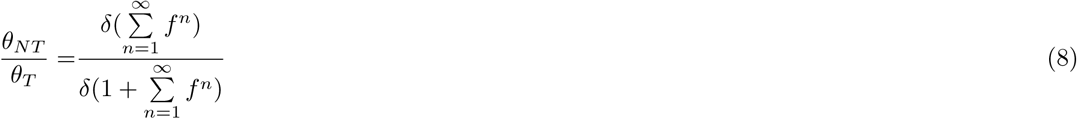

As shown, *δ* cancels out in the expected value of 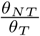. Therefore, the point estimates of *f* and *V*_*F*_ are influenced by the magnitude of VT and not by the predictive ability of the PGS (although the standard error of *V*_*F*_ increases as *δ* decreases; Kim et al., 2020, under review). Finally, because −1 *< f <* 1, the geometric series 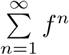 converges to 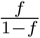, and thus

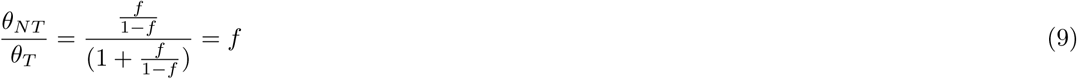

This demonstrates the same result in equation (6) again but from a different approach.

### Model 1: VT and AM

Model 1 assumes that the PGS explains all the trait heritability, as did Model 0, but now incorporates the influences of AM (Figure 2). As such, Model 1 yields estimates that are unbiased when there is AM, but only to the degree that the PGS captures the heritability of the trait. Given that the PGS’s for most traits explain little heritability (e.g., typically *<* 20%; Torkamani et al., 2018), the utility of this model is mostly didactic.

**Figure 2.**
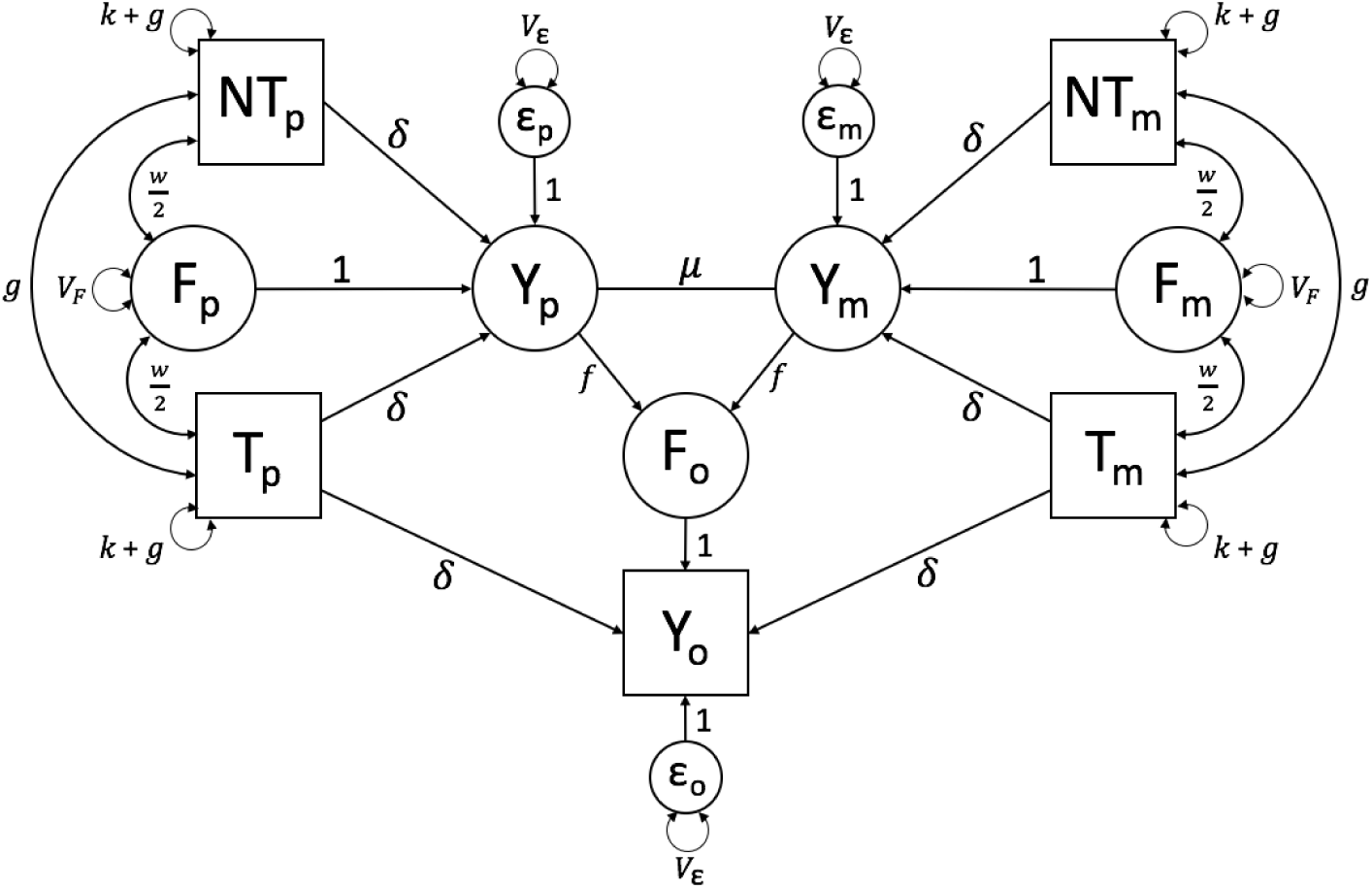
Path diagram of Model 1, which models the effects of AM and VT, assuming that the PGS explains the full trait heritability

We model AM using a copath, which follows a special set of path tracing rules, as explained above. The copath is represented as a straight line between *Y*_*p*_ and *Y*_*m*_ in Figure 2, and its path coefficient is denoted *µ*. The expected covariance between mates is all chains *Y*_*p*_ → *Y*_*m*_ or vice-versa. To traverse the copath, a chain must first be legitimate, so it must have already traversed a double-headed arrow. Thus, chains from *Y*_*p*_ → *Y*_*m*_ begin with the sum of all chains that connect *Y*_*p*_ back to itself (the sum of which = *V*_*Y*_, and each of which includes a double-headed arrow) before then crossing the copath (*µ*). At this point, the other path tracing rules reset, necessitating that each chain traverses another double-headed arrow. Thus, the chains end by traversing all chains from *Y*_*m*_ → *Y*_*m*_ (which also = *V*_*Y*_). Therefore, the covariance between mates is

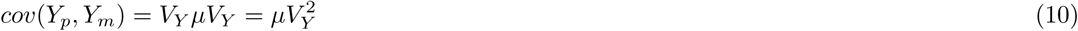

Note that 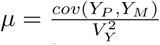 is neither the covariance 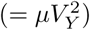 nor the correlation (= *µV*_*Y*_) between mates. AM and VT increase *V*_*Y*_ over time, and because we assume that the *correlation* between mates is constant across generations, the value of *µ* correspondingly decreases across generations until equilibrium is reached (which occurs in 5-10 generations). The information to estimate *µ* comes from the six observed covariances between haplotypic PGS’s as well as the four observed haplotypic PGS variances.

AM for a trait creates *gametic phase disequilibrium* between causal variants, meaning that trait-increasing alleles tend to coaggregate with other trait-increasing alleles and vice-versa. This occurs because similarity based on mates’ phenotypic scores implies similarity of genetic effects across mates as well. Two important consequences of gametic phase disequilibrium are the increase in genetic variation over what it would be in the absence of AM (in the base population), and the increase in genetic covariation between mates and close relatives (Lynch and Walsh, 1998).

A single generation of AM leads to covariance between the genetic scores of the maternal ([*N*]*T*_*m*_) and paternal ([*N*]*T*_*p*_) haplotypes, which is referred to as a “trans” covariance by Kong et al. and mediated by the copath in Model 1. However, two generations of AM (beginning in the grandparental generation) results in the recombination of alleles on the same haplotype, thus also leading to a “cis” covariance within the parental haplotypes. At equilibrium, after several generations of AM, the cis covariance (*cov*([*N*]*T*_∗_, [*N*]*T*_∗_)) equals the trans covariance (*cov*([*N*]*T*_*p*_, [*N*]*T*_*m*_)), with both denoted *g* in the models. Note, however, that only the cis covariances are explicit in Figure 2; the trans covariances are implicit, already being accounted for by *µ*. Note too that what is considered a trans covariance in the current generation (between *T*_*p*_ and *T*_*m*_) would be considered cis covariances in the next generation, when the offspring has children.

As denoted by the additional +*g* term in the haplotypic PGS variance in Figure 2, AM increases the variance within haplotypic PGS’s to the same degree as the covariance between them. The *k* term in the haplotypic PGS variance represents the variance of the haplotypic PGS in the base population, and is not estimated; rather, it is fixed depending on how the user scales the PGS. If the full PGS is standardized in the base population, then 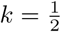. This value of *k* is useful because the increase in the variances of haplotypic genetic scores under AM is easily quantified by the degree to which it is greater than 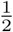. However, standardizing in the base population is typically impossible in real data, and so is mostly useful only in simulated data or when there is no AM (such as Model 0). In real data, the full PGS (*T*_*p*_ + *T*_*m*_) will typically be standardized in the current generation, in which case 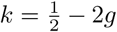. Finally, if the *haplotypic* PGS is scaled in the current generation to have a variance of 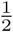, then 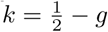. So long as the value of *k* is consistent with how the PGS is scaled, the estimates of other parameters will not be affected. In all cases, the variance of the full PGS (= *var*(*T*_*m*_ + *T*_*p*_) = 2*k* + 4*g*, which = 1 + 4*g* if the full PGS is standardized in the base population) = 1 if the full PGS is standardized in the current population, and = 1 + 2*g* if the haplotypic PGS is scaled to have variance 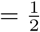 in the current population.

The increase in genetic (co)variance of the PGS under AM, *g*, can be obtained by constraining its value to all chains that connect [*N*]*T*_*p*_ → [*N*]*T*_*m*_ or vice-versa. Using Ω as a substitute for all chains [*N*]*T*_∗_ → *Y*_∗_,

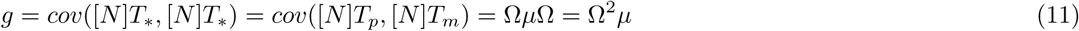

Of course, because of the additional variances and covariances between the haplotypic PGS’s, the expectation of Ω itself is different in this model than it was in Model 0. For Model 1, the expected value is:

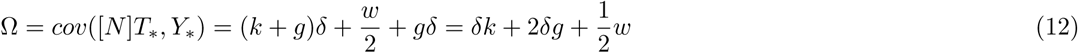

While accounting for AM makes this model more complicated than Model 0, substituting recurring chain segments drastically simplifies the derivations of parameters. For example, *θ*_*NT*_ — which is derived by counting all chains *Y*_*o*_ → *NT*_*p*_ and multiplying by 2 (to account for *Y*_*o*_ → *NT*_*m*_)— includes over 40 chains here as opposed to just 2 in Model 0. However, by using substitutions, this can be reduced to just four chains: 1) *Y*_*o*_ → *Y*_*p*_ → *NT*_*p*_ (= *f* Ω, the genetic nurture chain); 2) *Y*_*o*_ → *T*_*p*_ → *NT*_*p*_ (= *δg*, arising from the AM-induced covariance between *T*_*p*_ and *NT*_*p*_); 3) *Y*_*o*_ → *T*_*m*_ → *NT*_*p*_ (also = *δg*, arising from the AM-induced covariance between *T*_*m*_ and *NT*_*p*_); and 4) *Y*_*o*_ → *Y*_*m*_ → *Y*_*p*_ → *NT*_*p*_ (= *f V*_*Y*_ *µ*Ω, arising from the AM-induced covariance between *Y*_*m*_ and *NT*_*p*_). Therefore,

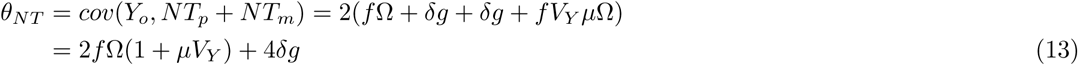

Similarly,

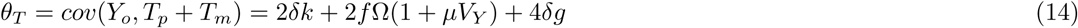

Thus, *θ*_*T*_ − *θ*_*NT*_ (= 2*δk*) is again an estimate of the direct effect of the PGS controlling for genetic nurture and, in this case, for AM.

In the same manner, the estimate of genetic nurture, *w*, can be derived by counting two chains *F*_*o*_ → *NT*_*m*_ and multiplying by 2 (to account for *F*_*o*_ → *NT*_*p*_): 1) *F*_*o*_ → *Y*_*m*_ → *NT*_*m*_ (= *f* Ω, the genetic nurture chain); and 2) *F*_*o*_ → *Y*_*p*_ → *Y*_*m*_ → *NT*_*m*_ (= *f V*_*Y*_ *µ*Ω, arising from the AM-induced covariance between *Y*_*p*_ and *NT*_*m*_). This leads to:

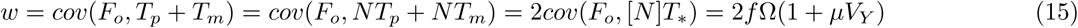

In Model 1, *w* remains an estimate of genetic nurture with its value being inflated by a factor (1 + *µV*_*Y*_) under AM. In the Kong et al. notation, direct genetic nurture (denoted *η*) refers to the aspect of *w* after removing the influence of AM, and thus *η* = 2*f* Ω. Kong et al. also denote *φ*_*η*_ as the added influence of AM on apparent genetic nurture, and thus *φ*_*η*_ = 2*f* Ω*µV*_*Y*_. We do not further make this distinction between direct and indirect genetic nurture. For completeness, it should be noted that *φ*_*δ*_ (the genetic covariance between *NT*_∗_ and *Y*_*o*_ induced by AM) in Kong et al.’s usage equals 4*δg* here. From this, it follows that *θ*_*NT*_ is no longer a direct estimate of genetic nurture (and that *θ*_*NT*_ ≠ *w*) when there is AM because some of the covariance between *Y*_*o*_ and *NT*_∗_ is now genetic in origin.

Finally, as was the case for *w*, the presence of AM causes the expectation of *V*_*F*_ to be inflated by a factor of (1 + *µV*_*Y*_):

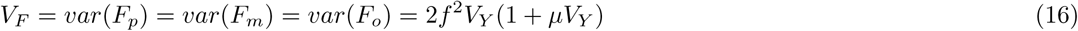

with the value of *V*_*Y*_ being similarly inflated in Model 1 versus Model 0:

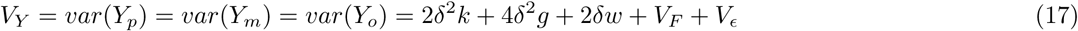

### Model 2: VT and AM with latent genetic effects

Model 2 builds on the concepts described above for modeling AM, but unlike Model 1, it provides unbiased estimates when there is AM and the PGS explains little trait heritability. It does this by modeling haplotypic latent genetic scores (LGS’s), denoted *L*[*N*]*T*_∗_ in Figure 3, that are defined to be statistically orthogonal to the haplotypic PGS’s ([*N*]*T*_∗_) in the base population. The latent genetic effects can be estimated either by including observed parental phenotypes, or by making an assumption about the base population additive genetic variance (and thus about the value of *a*). Here, we take the first approach by assuming that parental phenotypes are measured (hence the squares used to represent *Y*_*p*_ and *Y*_*m*_ in Figure 3), but discuss the second approach at the end of this section. It should be noted that full information maximum likelihood parameter estimates are unbiased by missingness unless the data is not missing at random (Schafer and Graham, 2002). Thus, all three phenotypes need not be measured in every family. Indeed, each family could be made up only of pairs (*Y*_*o*_, *Y*_*p*_; *Y*_*o*_, *Y*_*m*_; or *Y*_*m*_, *Y*_*p*_) and so long as all pairs are observed, parameter estimates would be unbiased, albeit with larger standard errors than with complete data.

**Figure 3.**
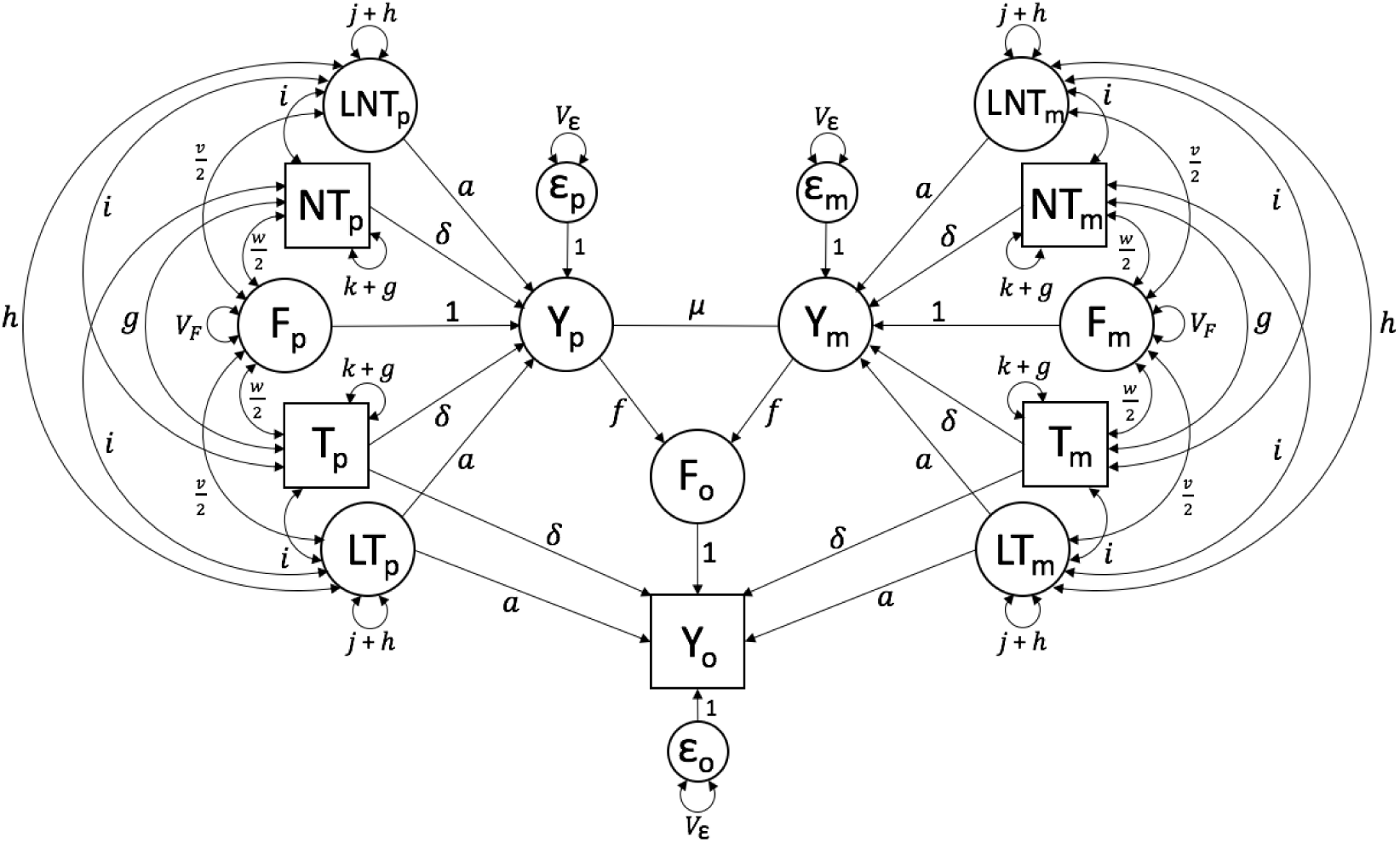
Path diagram of Model 2, which models the effects of AM, VT, and latent genetics

Model 2 includes two additional observed variables (*Y*_*p*_ and *Y*_*m*_), leading to a 7-by-7 observed variance-covariance matrix and 28 unique statistics. The four haplotypic PGS variances and the six covariances between them are used to estimate *g* and *µ*. Of the remaining 18 statistics, only six provide information that is not completely redundant to estimating parameters as specified in this model: the three described in Model 0 as well as the covariance between the parental phenotypes, the four covariances between one parent’s PGS and the other’s phenotype, and the two covariances between each parental phenotype with the offspring phenotype. The parent-offspring covariances are used to estimate the latent genetic path coefficient (*a*), which increases to the degree that *cov*(*Y*_∗_, *Y*_*o*_) is higher than expected after accounting for genetic covariance through *δ* and environmental covariance through *f*. In addition to *a*, there are four additional parameters (*j, h, v, i*) in this model. None of these are estimated. Rather, their values are determined from non-linear constraints, which we turn to in order.

The variance of the haplotypic LGS (= *j* + *h*) is treated analogously to the variance of the haplotypic PGS (=*k* + *g*). Like *k, j* is defined as the genetic variance of the haplotypic LGS in the base population; however, unlike *k* (which is measured and therefore depends upon how the PGS is scaled), *j* is the variance of a latent construct and could thereby take any arbitrary value. The simplest choice is to define *j* so as to be consistent with *k*. Specifically, if the PGS is standardized in the base population (where 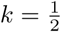), then 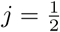. If the PGS is standardized at equilibrium to have a variance of 1 (where 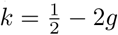, then 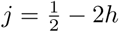. If the haplotypic PGS is scaled at equilibrium to have a variance of 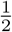 (where 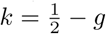.), then 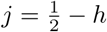.

The increase in the variance of the haplotypic LGS due to AM (*h*) can be estimated under the reasonable assumption that the increase in the variance of the LGS from the base to the current population is proportionate to that of the PGS from the base to the current population. This assumption could be violated if the genes that drive the PGS association are more or less correlated with the trait upon which AM is actually occurring than the genes underlying the LGS, which seems unlikely. This assumption is equivalent to 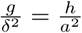, which leads to

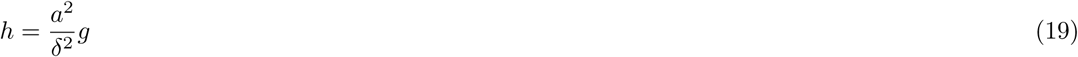

Thus, *h* and *g* are the same only when the PGS and LGS explain the same amount (half) of the total heritability. Furthermore, similar to Ω, we define Γ to be the covariance between a parent’s phenotype and one of their LGS’s (Γ = *cov*(*L*[*N*]*T*_∗_, *Y*_∗_). Using Γ as a shortcut, the expected value of *h* can also be found by path tracing

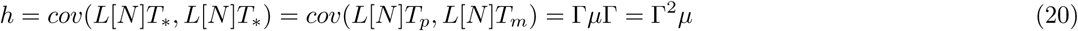

Setting these two values of *h* to be equal leads to the nonlinear constraint

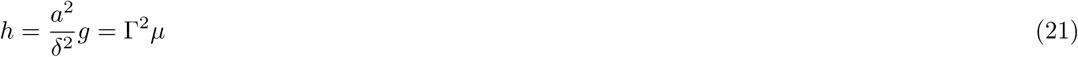

To enable estimation of the covariance between *F* and the LGS’s (denoted by *v*), we make a similar assumption that the ratio of genetic nurture to direct genetic effects is the same for observed as for latent genetic effects. This assumption could be violated if the genes driving the PGS association are more or less correlated with the trait that VT works through than the genes underlying the LGS, which again seems unlikely. This assumption is equivalent to 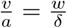, which leads to

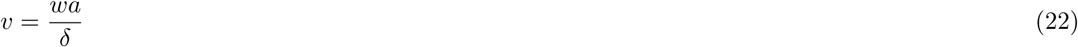

The expected value of *v* via path tracing, and the resulting non-linear constraint, are

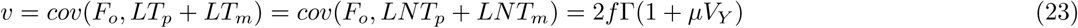

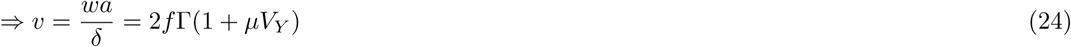

For the same reason that AM induces a covariance among PGS’s (*g*) and among LGS’s (*h*), it also induces a covariance between PGS’s and LGS’s, which we call *i*. From path tracing, the expected value of *i* is

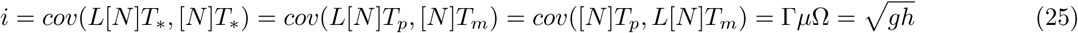

Unlike Model 1, Model 2 yields unbiased estimates of the full *V*_*F*_, genetic nurture, and additive genetic variation, even when there is AM and the PGS explains only a fraction of total heritability. Model 2 properly accounts for the additional covariance between the PGS and the offspring phenotype that arises from the AM-induced covariance between PGS and LGS. For example,

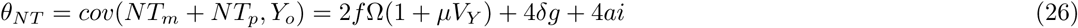

where

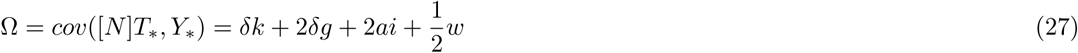

Thus, the PGS-LGS covariance (*i*) inflates both the genetic nurture part of *θ*_*NT*_ (2*f* Ω(1 + *µV*_*Y*_)) as well as the genetic part that arises via AM (4*δg* + 4*ai*). When the PGS explains a small portion of the heritability, the covariance between the LGS and the PGS can be much greater than the covariance between the PGS’s themselves (*i* >> *g*). By not accounting for *i*, the observed *θ*_*NT*_ is inflated over its expected value in Model 1, upwardly biasing estimates of *V*_*F*_ and genetic nurture (Kim et al., 2020, under review).

Several other parameters in this model are the latent genetic analogs to parameters related to the observed PGS’s, including *θ*_*LNT*_ (= *cov*(*Y*_*o*_, *LNT*_*p*_ + *LNT*_*m*_), the analog to *θ*_*NT*_) and *θ*_*LT*_ (= *cov*(*Y*_*o*_, *LT*_*p*_ + *LT*_*m*_), the analog to *θ*_*T*_). Expectations of these and other parameters that have not been derived in this section can be found in the Supplement.

Finally, as noted above, Model 2 can be fit without using parental phenotypes if there exist good estimates of the total heritability in the base population. For a standardized trait, the additive genetic variation in the base population is *a*^2^ + *δ*^2^; thus, by subtraction of the *δ*^2^ term observed in the data (where *δ* = *θ*_*T*_ − *θ*_*NT*_), one can find the assumed value of *a* and substitute it into the model. This leads to unbiased estimates of all parameters whenever the assumed value of *a* is correct and downwardly biased estimates of *w* and *V*_*F*_ to the degree that the assumed value of *a* is too high (and vice-versa). When using this strategy, it is therefore important to have decent estimates of heritability in the base population that account for the influences of genetic nurture and AM. Estimates from twin studies are biased under these conditions, but estimates from extended twin family models should be much less so (Keller et al., 2010). Kong et al. recognized the confounding influence that the LGS has on parameter estimates when there is AM, and attempted to estimate genetic nurture using assumed values of the base population heritability that came from relatedness disequilibrium regression (RDR; Young et al., 2018). However, estimates of heritability from RDR are actually estimates of the base population genetic variance scaled by the phenotypic variance in the current population, and are therefore biased downwards under AM (Kemper et al., in prep.). This likely led to overestimates of genetic nurture in Kong et al., and would have led to overestimates of *V*_*F*_ had it been estimated. Nonetheless, if the equilibrium spousal correlations are known, a simple correction can be applied to RDR estimates of heritability (Kemper et al., in prep.).

### Accounting for differences in parent vs. offspring phenotypes

In all of these models, we have assumed that the genetic (PGS and LGS) effects are equivalent in parents and offspring. This assumption would be violated if there are gene-by-age or gene-by-cohort effects. Kong et al. provide some evidence of this in their data: the correlation between the PGS and educational attainment was significantly higher among offspring (.22) than among parents (.12). When parental phenotypes are measured in Model 2, accounting for such effects is possible by estimating two different *δ* values, one for offspring (*δ*_*o*_) and one for parents (*δ*_∗_). While information for estimating *δ*_*o*_ still comes directly from *θ*_*T*_ − *θ*_*NT*_, there is no direct estimate of *δ*_∗_, even though it is informed by *cov*(*Y*_∗_, *T*_∗_ + *NT*_∗_). A reasonable assumption, such as equal proportions of direct genetic effects 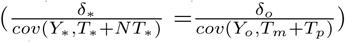, should allow estimation of both *δ*_∗_ and *δ*_*o*_, making this model identified, although we have not checked this formally. On the other hand, when *cov*(*Y*_∗_, *T*_∗_ + *NT*_∗_) is not observed, inter-generational differences could be modeled by fixing *δ*_∗_ if there are independent estimates of the PGS effects in the parental generation.

It may also be of interest to understand how one parental trait influences a different offspring trait. For example, Kong et al. showed a covariance between the nontransmitted PGS of educational attainment and offspring health, consistent with cross-trait (parental education to offspring health) VT and genetic nurture. Such cross-trait genetic nurture would contribute to apparent genetic correlations that have nothing to do with pleiotropy. To investigate cross-trait VT using the above example under the current framework, one could use the PGS and parental values of educational attainment along with the offspring values of health, and plug them into the above models without modification. This is similar to the approach taken by Kong et al. However, such an approach does not account for AM within (health - health) or across (education - health) traits, nor does it account for within-trait genetic nurture effects. For these reasons, we believe that cross-trait VT and genetic nurture effects are optimally modeled bivariately, using the PGS’s and phenotypic values of the two traits in both parents and offspring; including more than two traits would also be possible, but results from such a model would likely be incomprehensible. The parameters from the above models would be 2-by-2 full (in the case of path coefficients) or symmetric (in the case of variance - covariance) matrices. Conveniently, nothing about the derivations in this paper would change except for keeping track of when matrices should be transposed, which obeys an additional path analysis rule (Vogler and Cockerham, 1985). This bivariate model would estimate two direct and two cross-trait VT paths and four genetic nurture paths all while accounting for direct genetic effects, pleiotropy, and AM within and between traits. While this sounds like a lot to ask of a model, there is a tremendous amount of unique information contained in the 14-by-14 observed variance-covariance matrix, making this approach powerful if the PGS *r*^2^’s for both traits are nontrivial. Development of bivariate extensions of the present models is left for future work.

### Testing and modeling different mechanisms of AM

While our models have thus far assumed primary phenotypic AM under equilibrium, other mechanisms of assortment are possible. Indeed, there is considerable power for testing different mechanisms of AM, which could itself be a focus of these models. This power stems from the amount of information in this model relevant to AM. There are a total of 10 observed haplotypic PGS variances or covariances which collectively provide 10 estimates of *g* (see Model 1). The *χ*(1) test of whether the average value across all 10 *g* estimates is significantly greater than 0 tests whether mate similarity (either on the trait in question or a trait genetically correlated to it) has led to genetic covariance, as predicted by primary AM. Additionally, of these 10 estimates of *g*, 6 provide information on cis (within-person) genetic covariances and 4 provide information on trans (across-mate) genetic covariances. The *χ*(1) test of whether these two groups of covariances are equal tests whether AM has gone on long enough to lead to equilibrium values of parameters. Significantly higher estimates of trans *g* vs. cis *g* would suggest that AM is at disequilibrium. Similarly, if trans *g* estimates are significantly greater than 0 while cis *g* estimates are not, this would support the hypothesis that only a single generation of AM has occurred (which is consistent with what Kong et al. found for educational attainment in Iceland). Furthermore, the ten estimates of *g* can be used to derive expected values of *cov*(*NT*_*p*_ + *T*_*p*_, *Y*_*m*_), *cov*(*NT*_*m*_ + *T*_*m*_, *Y*_*p*_), and *cov*(*Y*_*p*_, *Y*_*m*_) to test various models of AM. For example, mate similarity caused by environmental similarity (social homogamy) predicts that observed *cov*(*Y*_*p*_, *Y*_*m*_) is higher than that implied by *g*. On the other hand, primary AM on a trait that is more genetically than phenotypically correlated with the measured trait (a form of genetic homogamy) would predict that observed *cov*(*Y*_*p*_, *Y*_*m*_) is lower than that implied by *g*

Once the data suggest a particular mechanism underlying mate similarity, it can be modeled using the present framework. For example, in the Supplement, we show how disequilibrium AM from a single generation of AM can be modeled by setting expectations of cis *g, h*, and *i* to zero. Furthermore, social and genetic homogamy can be modeled by assuming that AM occurs on a latent trait, 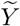, that is related to *Y* through either environmental or genetic pathways, respectively (Keller et al., 2009). This allows the observed *cov*(*Y*_*p*_, *Y*_*m*_) to differ from the *cov*(*Y*_*p*_, *Y*_*m*_) implied by *g*. Thus, alternative mechanisms of AM can be formally tested using the rich information available from parent and offspring PGS’s and phenotypes, and when called for, models can be modified to incorporate alternative mechanisms of AM.

## Discussion

Estimates of genetic effects from GWAS or heritability studies do not necessarily reflect the biological action of genes; they can be amplified by genetic nurture, their covariance with environmental influences provided by the parents. While estimating direct genetic effects after removing their covariance with the environment is important, we argue that the converse—estimating the direct environmental effect after removing its covariance with genetic effects—is at least as important. The models presented above allow for estimates of this direct environmental influences of parents on offspring, and also suggest several important extensions. For example, there exists much more GWAS data with siblings than with parents, and so extending the models above to include siblings, twins, and potentially other collateral relatives remains a next step. Furthermore, as shown by Kong et al., with enough data it is straightforward to estimate differential genetic nurture effects depending on the parent of origin; thus, it is equally straightforward to estimate differential VT effects from fathers vs. mothers. Finally, as noted above, multivariate extensions to this model would provide insight into how one parental trait influences a different childhood trait, which serves as an additional avenue for future work.

There are several important caveats and limitations to the present approach. We discuss here only the central ones. One very important caveat is that estimated *V*_*F*_ from these models cannot be considered the full parental effect on this trait. Rather, it is the variance in the trait due to parental influences that are associated with *the specific trait assessed by the PGS*. For example, the vertical transmission variance in a model that uses an educational attainment PGS only estimates how traits genetically related to parental educational attainment influence offspring educational attainment. If other parental traits, such as intelligence, warmth, work ethic, conscientiousness, etc. also influence offspring educational attainment, then the portion of variance due to these and other parental influences that are genetically uncorrelated to educational attainment will be missed.

The above caveat is related to the limitation that, in order to accurately characterize the influence of a parental trait on an offspring trait, sufficiently predictive PGS’s (e.g., *r*^2^ > .02) must exist for the traits most relevant to parenting. Optimally (to use Model 2), these traits should also be measured in parents in the same dataset that the models are applied to. This is, perhaps, the greatest weakness of the current approach: it can only look under the lamppost, at traits analyzed in large GWAS’s for which sufficiently predictive PGS’s exist. Because it is so easily and frequently collected, educational attainment may be an exception, but a great many traits relevant to parental influences have never been investigated in GWAS. This—the ability to use PGS’s to understand how parents influence offspring—is another motivation to continue to extend GWAS investigations from their traditional focus on health to as many behavioral and psychological traits as possible.

That said, there are many traits that have sufficiently predictive PGS’s to answer questions of great interest. For example, does parental subjective well being, liability to major depression, schizophrenia, or externalizing disorder directly influence the same or different traits in the children (Okbay et al., 2016; Barr et al., 2020)? Does parental socioeconomic status directly impact offspring socioeconomic status, educational attainment, or subjective well being (Hill et al., 2019)? Does parental smoking influence offspring smoking (Liu et al., 2019)? This latter question is interesting with respect to negative VT, which occurs when higher values of the parental trait decrease the offspring trait. Negative VT would lead to positive *V*_*F*_ but to negative genetic nurture, dampening estimated genetic influences from GWAS or heritability studies. While this is probably rare, there is some evidence from extended twin family models that negative VT occurs for smoking (Maes et al., 2006). While this finding has been explained away in the literature as probably being driven by gene-by-age interactions, it is also possible that smoking and other traits associated with teen rebelliousness lose their lustre when parents engage in them. Given that a sufficiently predictive PGS for smoking behaviors exists (Liu et al., 2019) and that Model 2 can be extended to account for gene-by-age effects, a whole-genome dataset that includes parents, offspring, and information on smoking behavior could resolve whether parental smoking directly increases or decreases offspring smoking.

A further caveat to the above approach is that stratification can bias estimates if it is not properly controlled for. For example, if the discovery GWAS for educational attainment did not fully correct for mean differences across ancestry groups, the PGS for educational attainment would predict both educational attainment as well as ancestry. In the models explored above, this stratification effect would increase the covariance between the transmitted and nontransmitted PGS’s and the offspring phenotype (*θ*_[*N*]*T*_), inflating evidence of genetic nurture and *V*_*F*_. While stratification is a type of passive G-E covariance that inflates parent-offspring resemblance, the mechanism is due to a factor (ancestry) that is shared between parents and offspring rather than a direct parental-to-offspring influence, and so these should be disambiguated. Therefore, to minimize the effects of stratification, principal components from the genomic relationship matrices should be included as covariates in both the discovery GWAS as well as the causal models discussed above.

Lastly, we have assumed that the passive G-E covariance we model arises only from VT (genetic nurture) as opposed to horizontal (such as sibling) transmission. For certain traits, such as experimentation with drugs and alcohol, horizontal transmission seems at least as likely as VT. In a model that includes both parents and siblings, there would be sufficient power to differentiate horizontal transmission from VT. In the meantime, it must be kept in mind that estimates of *V*_*F*_ from these models may also contain environmental influences from siblings or (less likely) from other collateral relatives.

To our knowledge, this is the first treatment of how transmitted and nontransmitted PGS’s can be used to estimate the direct effect of parents on their offspring. It builds upon the seminal work by Kong et al., who recognized that this data could be used to estimate genetic nurture. There has been long-standing interest in how parents influence offspring in fields such as developmental psychology, but the traditional approach of correlating parental behavior with offspring outcomes does not control for the confounding influence of shared genetics. Given the skepticism of genetic approaches in fields dedicated to studying parenting, it is perhaps ironic that molecular genetic data provides an excellent way to estimate the direct effect of parents on offspring. Genome-wide data, originally collected to find the specific alleles that underlie health-related traits, has begun to be used for a multitude of purposes never envisioned by early practitioners. As we have argued here, another one of these purposes is to help disentangle how parents influence their children—genetically, environmentally, and in concert.

## Acknowledgments

We thank Richard Border, Hermine Maes, and David Evans for help with ideas and feedback as the project developed. This work was supported by grants R01MH100141 (to MCK) and T32MH016880 (to Dr. John Hewitt) and the Institute for Behavioral Genetics. This work utilized resources from the University of Colorado Boulder Research Computing Group, which is supported by the National Science Foundation (awards ACI-1532235 and ACI-1532236), the University of Colorado Boulder, and Colorado State University.

## Notes

### Competing Interest Statement

The authors have declared no competing interest.

