## Supplementary Material for "Estimation of parental effects using polygenic scores"

Estimation of parental effects using polygenic scores: Supplementary Material

### 1 Path Diagrams

#### 1.1 No Assortative Mating, Equilibrium (Observed PRS Only)

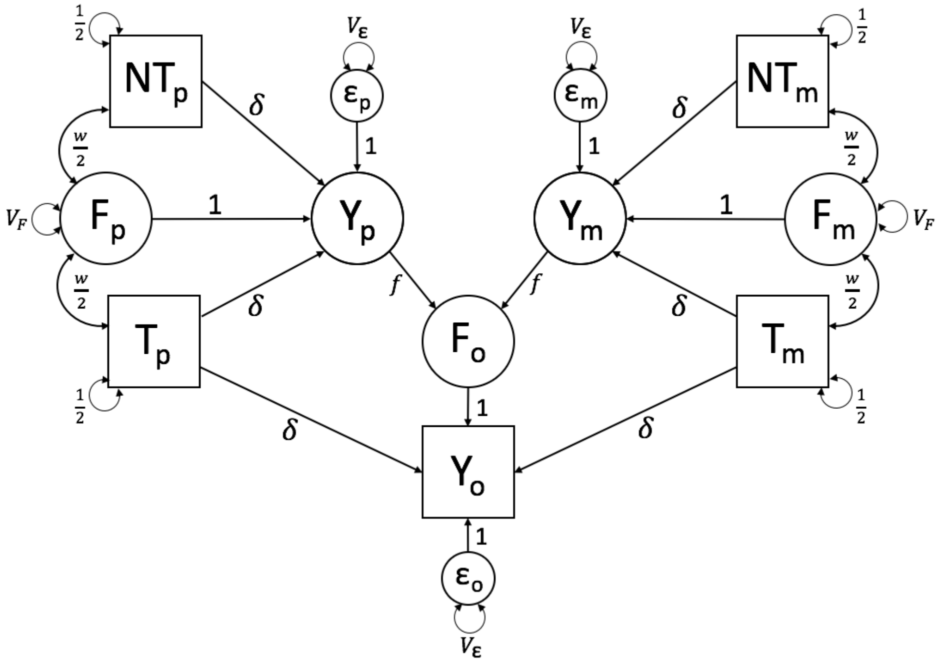


**Supp. Figure 1.** Path diagram modeling the effects of VT at equilibrium, assuming that the PGS explains the full trait heritability and that there is no AM.

#### 1.2 No Assortative Mating, Disequilibrium (Observed PRS Only)

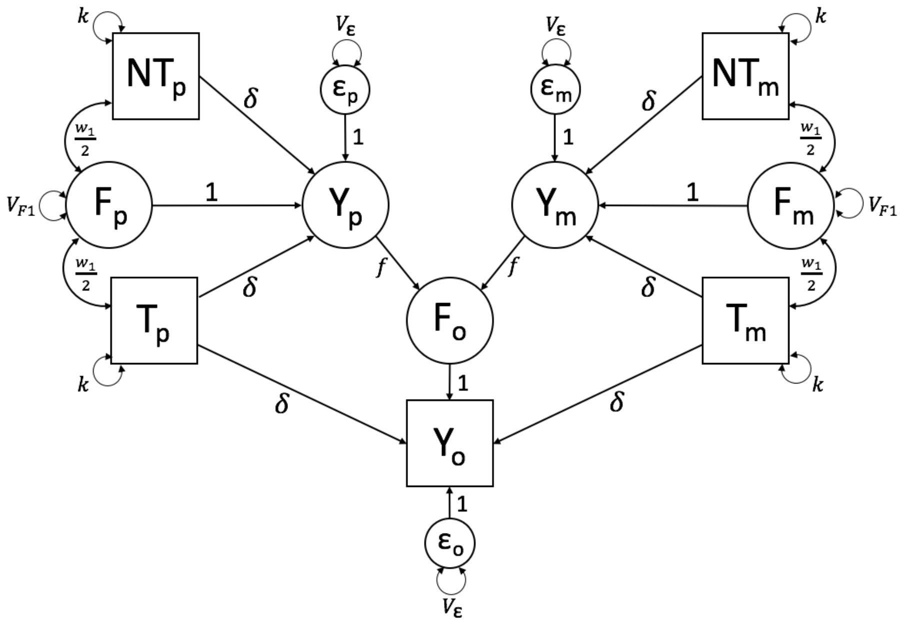


**Supp. Figure 2.** Path diagram modeling the effects of VT at disequilibrium, assuming that the PGS explains the full trait heritability and that there is no AM. Because the model is at disequilibrium, subscripts 1 (shown) and 2 (implied) are necessary to differentiate constraints in the parental generation from constraints in the offspring generation.

#### 1.3 Assortative Mating, Equilibrium (Observed PRS Only)

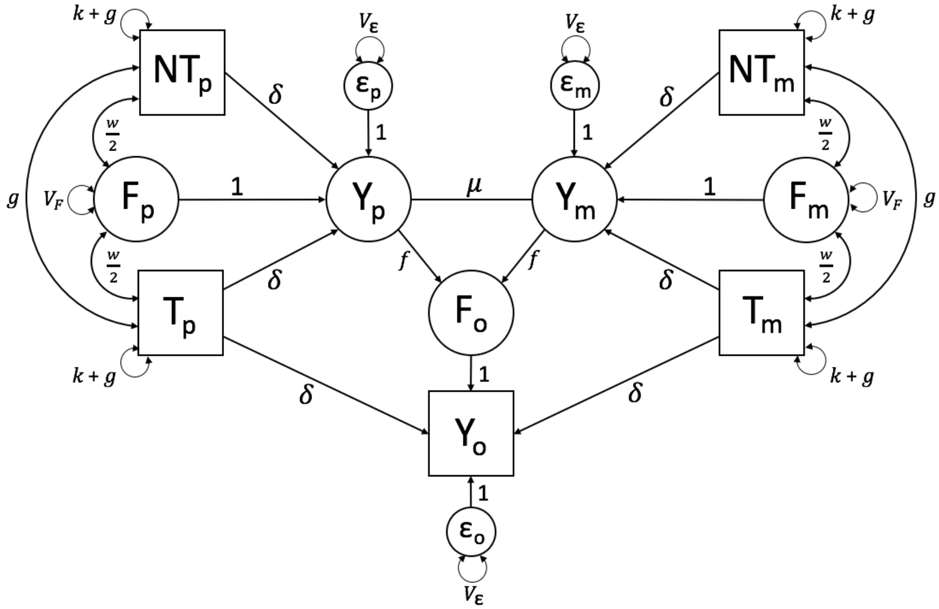


**Supp. Figure 3.** Path diagram modeling the effects of VT and AM at equilibrium, assuming that the PGS explains the full trait heritability.

#### 1.4 Assortative Mating, Disequilibrium (Observed PRS Only)

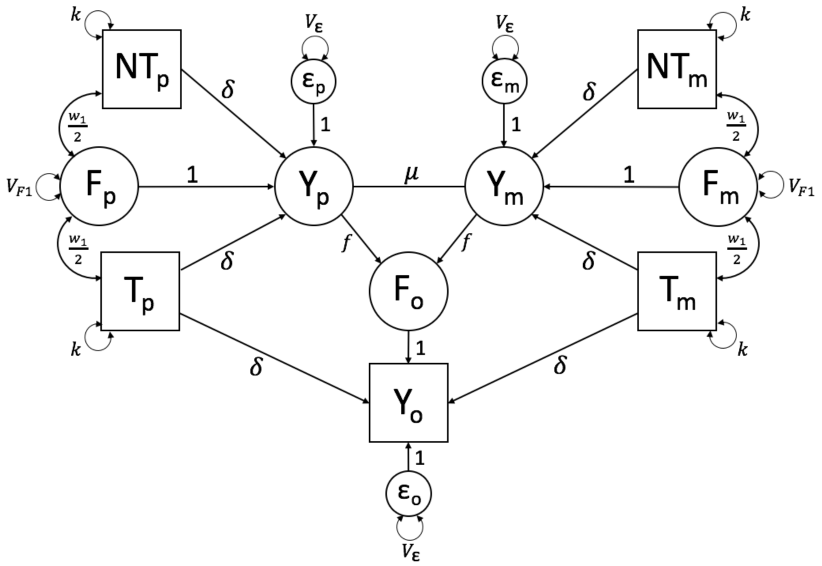


**Supp. Figure 4.** Path diagram modeling the effects of VT and AM at disequilibrium (after one generation of AM), assuming that the PGS explains the full trait heritability. Because the model is at disequilibrium, subscripts 1 (shown) and 2 (implied) are necessary to differentiate constraints in the parental generation from constraints in the offspring generation.

#### 1.5 No Assortative Mating, Equilibrium (Latent and Observed PRS)

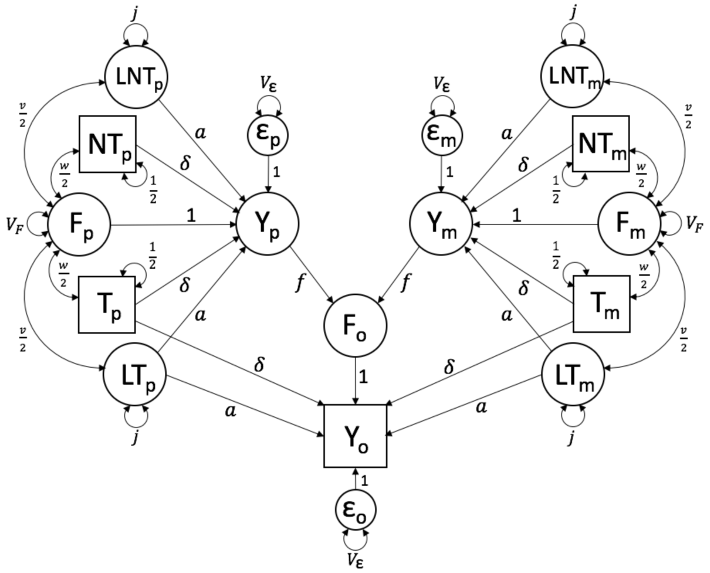


**Supp. Figure 5.** Path diagram modeling the effects of VT at equilibrium, assuming that there is no AM. Because $Y_p$ and $Y_m$ are unobserved in this Model (hence the circles used to represent them), the value of $a$ must be assumed prior to running the model; otherwise, the model will be under-identified. Alternatively, one could use observed parental phenotypes, thereby allowing them to estimate the value of $a$.

#### 1.6 No Assortative Mating, Disequilibrium (Latent and Observed PRS)

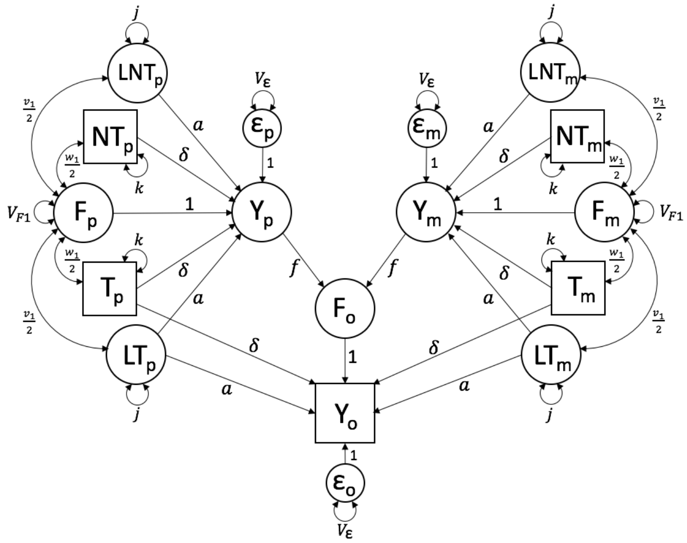


**Supp. Figure 6.** Path diagram modeling the effects of VT at disequilibrium, assuming no AM. Because $Y_p$ and $Y_m$ are unobserved in this Model (hence the circles used to represent them), the value of $a$ must be assumed prior to running the model; otherwise, the model will be under-identified. Alternatively, one could use observed parental phenotypes, thereby allowing them to estimate the value of $a$. Also, because the model is at disequilibrium, subscripts 1 (shown) and 2 (implied) are necessary to differentiate constraints in the parental generation from constraints in the offspring generation.

#### 1.7 Assortative Mating, Equilibrium (Latent and Observed PRS)

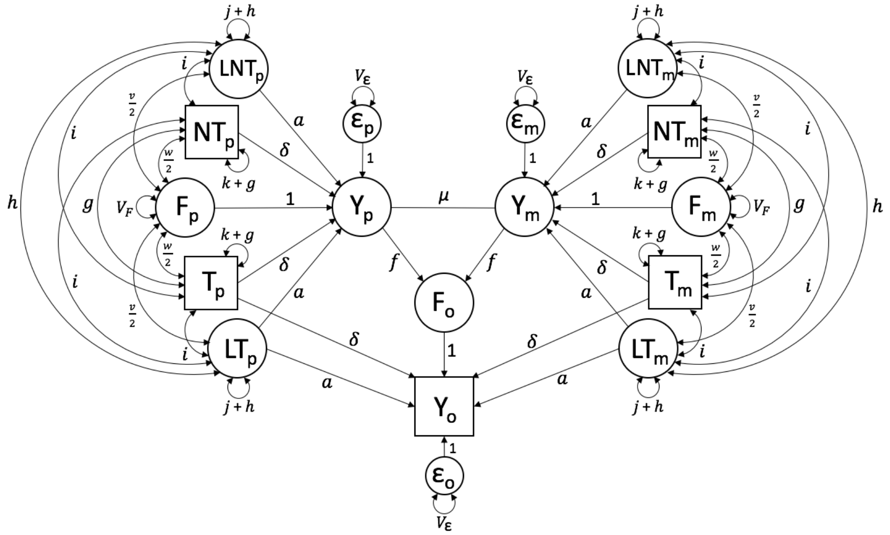


**Supp. Figure 7.** Path diagram modeling the effects of VT and AM at equilibrium. Because $Y_p$ and $Y_m$ are unobserved in this Model (hence the circles used to represent them), the value of $a$ must be assumed prior to running the model; otherwise, the model will be under-identified. Alternatively, one could use observed parental phenotypes, thereby allowing them to estimate the value of $a$.

#### 1.8 Assortative Mating, Disequilibrium (Latent and Observed PRS)

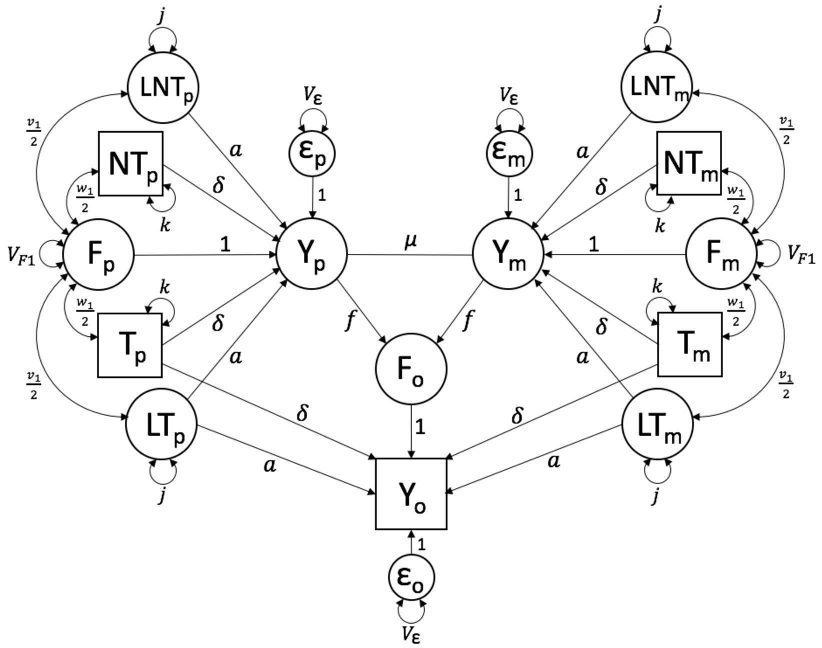


**Supp. Figure 8.** Path diagram modeling the effects of VT and AM at disequilibrium (after one generation of AM). Because $Y_p$ and $Y_m$ are unobserved in this Model (hence the circles used to represent them), the value of $a$ must be assumed prior to running the model; otherwise, the model will be under-identified. Alternatively, one could use observed parental phenotypes, thereby allowing them to estimate the value of $a$. Also, because the model is at disequilibrium, subscripts 1 (shown) and 2 (implied) are necessary to differentiate constraints in the parental generation from constraints in the offspring generation.

**Supp. Table I: Equilibrium Model Constraints**

| | **No AM, No Latent PGS's** | **No AM, Latent PGS's** | **AM, No Latent PGS's** | **AM, Latent PGS's** |
|---|---|---|---|---|
| $\Omega$ | $\frac{1}{2}(\delta + w)$ | $\frac{1}{2}(\delta + w)$ | $\delta k + \frac{1}{2}w + 2\delta g$ | $\delta k + \frac{1}{2}w + 2\delta g + 2ai$ |
| $g$ | 0 | 0 | $\Omega^2\mu$ | $\Omega^2\mu$ |
| $V_F$ | $2f^2V_Y$ | $2f^2V_Y$ | $2f^2V_Y(1 + V_Y\mu)$ | $2f^2V_Y(1 + V_Y\mu)$ |
| $w$ | $2f\Omega$ | $2f\Omega$ | $2f\Omega(1 + V_Y\mu)$ | $2f\Omega(1 + V_Y\mu)$ |
| $\theta_{NT}$ | $2f\Omega$ =w | $2f\Omega$ = w | $4\delta g + 2w$ | $4\delta g + 4ai + 2w$ |
| $\theta_T$ | $\delta + 2f\Omega$ $= \delta + \theta_{NT}$ | $\delta + 2f\Omega$ $= \delta + \theta_{NT}$ | $2\delta k + 4\delta g + 2w$ $= 2\delta k + \theta_{NT}$ | $2\delta k + 4\delta g + 4ai + 2w$ $= 2\delta k + \theta_{NT}$ |
| $V_Y$ | $2\delta\Omega + \delta w + V_F + V_\epsilon$ | $2a\Gamma + 2\delta\Omega + av$ $+ \delta w + V_F + V_\epsilon$ | $2\delta\Omega + \delta w + V_F + V_\epsilon$ | $2a\Gamma + 2\delta\Omega + av$ $+ \delta w + V_F + V_\epsilon$ |
| $\Gamma$ | – | $aj + \frac{1}{2}v$ | – | $aj + \frac{1}{2}v + 2ah + 2\delta i$ |
| $v$ | – | $2f\Gamma$ | – | $2f\Gamma(1 + V_Y\mu)$ |
| $\theta_{LNT}$ | – | $2f\Gamma$ = v | – | $4ah + 4\delta i + 2v$ |
| $\theta_{LT}$ | – | $2aj + v$ = 2aj + $\theta_{LNT}$ | – | $2aj + 4ah + 4\delta i + 2v$ = 2aj + $\theta_{LNT}$ |
| $h$ | – | 0 | – | $\Gamma^2\mu$ |
| $i$ | – | 0 | – | $\Omega\mu\Gamma$ $= \sqrt{g_2h_2}$ |
| cov($Y_p$, $L[N]T_m$) cov($Y_m$, $L[N]T_p$) | – | 0 | – | $V_Y\mu\Gamma$ |
| cov($Y_p$, $[N]T_m$) cov($Y_m$, $[N]T_p$) | 0 | 0 | $V_Y\mu\Omega$ | $V_Y\mu\Omega$ |
| cov($Y_p$, $F_m$) cov($Y_m$, $F_p$) | 0 | 0 | $V_Y\mu(\delta w + V_F)$ | $V_Y\mu(\delta w + av + V_F)$ |
| cov($F_p$, F$_m$) | 0 | 0 | $\mu(\delta w + V_F)^2$ | $\mu(\delta w + av + V_F)^2$ |
| cov($Y_p$, $Y_m$) | 0 | 0 | $V_Y^2\mu$ | $V_Y^2\mu$ |
| cov($Y_o$, $Y_*$) | $\delta\Omega + fV_Y$ | $\delta\Omega + a\Gamma + fV_Y$ | $(\delta\Omega + fV_Y)(1 + V_Y\mu)$ | $(\delta\Omega + a\Gamma + fV_Y)(1 + V_Y\mu)$ |
| cov($Y_*$, $F_o$) | $fV_Y$ | $fV_Y$ | $fV_Y(1 + V_Y\mu)$ | $fV_Y(1 + V_Y\mu)$ |
| cov($Y_*$, $F_*$) | $\delta w + V_F$ | $\delta w + av + V_F$ | $\delta w + V_F$ | $\delta w + av + V_F$ |

Parameter constraints under equilibrium. Here, we use the $*$ subscript to denote either $p$ or $m$ but not both within a single term. For example, $cov(Y_*, [N]T_*)$ can be written in the place of $cov(Y_p, T_p)$, $cov(Y_p, NT_p)$, $cov(Y_m, T_m)$, or $cov(Y_m, NT_m)$. However, $cov(Y_*, T_*)$ does not equal $cov(Y_p, T_m)$ or $cov(Y_m, T_p)$, which mix $m$ and $p$ in the same term.

Supp. Table II: Disequilibrium Model Constraints

| | **No AM, No Latent PGS's** | **No AM, Latent PGS's** | **AM, No Latent PGS's** | **AM, Latent PGS's** |
|---|---|---|---|---|
| $\Omega_1$ | $\frac{1}{2}(\delta + w_1)$ | $\frac{1}{2}(\delta + w_1)$ | $\delta k + \frac{1}{2}w_1$ | $\delta k + \frac{1}{2}w_1$ |
| $\Omega_2$ | $\frac{1}{2}(\delta + w_2)$ | $\frac{1}{2}(\delta + w_2)$ | $\delta k + \frac{1}{2}w_2 + \delta g$ | $\delta k + 2ai_2 + \frac{1}{2}w_2 + \delta g_2$ |
| $g_1$ | 0 | 0 | 0 | 0 |
| $g_2$ | 0 | 0 | $\Omega_1^2\mu$ | $\Omega_1^2\mu$ |
| $V_{F1}$ | $2f^2V_{Y1}$ | $2f^2V_{Y1}$ | $2f^2V_{Y1}$ | $2f^2V_{Y1}$ |
| $V_{F2}$ | $2f^2V_{Y2}$ | $2f^2V_{Y2}$ | $2f^2V_{Y1}(1+V_{Y1})$ | $2f^2V_{Y1}(1+V_{Y1})$ |
| $w_1$ | $2f\Omega_1$ | $2f\Omega_1$ | $2\Omega_1\mu(V_{F1} + \delta w_1)$ | $2\Omega_1\mu(V_{F1} + \delta w_1 + av_1)$ |
| $w_2$ | $2f\Omega_2$ | $2f\Omega_2$ | $2f\Omega_1(1 + V_{Y1}\mu)$ | $2f\Omega_1(1 + V_{Y1}\mu)$ |
| $\theta_{NT}$ | $2f\Omega_1$<br>$= \mathrm{w}_1$ | $2f\Omega_1$<br>$= \mathrm{w}_1$ | $2w_2 + 2\delta g_2$ | $w_2 + 2\delta g_2 + 2ai_2$ |
| $\theta_T$ | $\delta + 2f\Omega_1$<br>$= \delta + \theta_{NT}$ | $\delta + 2f\Omega_1$<br>$= \delta + \theta_{NT}$ | $2\delta k + w_2 + 2\delta g_2$<br>$= 2\delta k + \theta_{NT}$ | $2\delta k + w_2 + 2\delta g_2 + 2ai_2$<br>$= 2\delta k + \theta_{NT}$ |
| $V_{Y1}$ | $2\delta\Omega_1 + \delta w_1$<br>$+ \mathrm{V}_{F1} + V_\epsilon$ | $2\delta\Omega_1 + 2a\Gamma_1 + \delta w_1$<br>$+ \mathrm{av}_1 + V_{F1} + V_\epsilon$ | $2\delta\Omega_1 + \delta w_1 + V_{F1} + V_\epsilon$ | $2\delta\Omega_1 + 2a\Gamma_1 + \delta w_1$<br>$+ \mathrm{av}_1 + V_{F1} + V_\epsilon$ |
| $V_{Y2}$ | $2\delta\Omega_2 + \delta w_2$<br>$+ \mathrm{V}_{F2} + V_\epsilon$ | $2\delta\Omega_2 + 2a\Gamma_2 + \delta w_2$<br>$+ \mathrm{av}_2 + V_{F2} + V_\epsilon$ | $2\delta\Omega_2 + \delta w_2 + V_{F2} + V_\epsilon$ | $2\delta\Omega_2 + 2a\Gamma_2 + \delta w_2$<br>$+ \mathrm{av}_2 + V_{F2} + V_\epsilon$ |
| $\Gamma_1$ | – | $aj + \frac{1}{2}v_1$ | – | $aj + \frac{1}{2}v_1$ |
| $\Gamma_2$ | – | $aj + \frac{1}{2}v_2$ | – | $aj + 2\delta i_2 + \frac{1}{2}v_2 + ah_2$ |
| $v_1$ | – | $2f\Gamma_1$ | – | $2\Gamma_1\mu(V_{F1} + \delta w_1 + av_1)$ |
| $v_2$ | – | $2f\Gamma_2$ | – | $2f\Gamma_1(1 + V_{Y1}\mu)$ |
| $\theta_{LNT}$ | – | $2f\Gamma_1$<br>$=\mathrm{v}_1$ | – | $v_2 + 2\delta i_2 + 2ah_2$ |
| $\theta_{LT}$ | – | $2f\Gamma_2$ | – | $2aj + v_2 + 2\delta i_2 + 2ah_2$<br>$= 2\ \mathrm{aj} + \theta_{LNT}$ |
| $h_1$ | – | 0 | – | 0 |
| $h_2$ | – | 0 | – | $\Gamma_1^2\mu$ |
| $i_1$ | – | 0 | – | 0 |
| $i_2$ | – | 0 | – | $\Omega\mu\Gamma$<br>$= \sqrt{g_2 h_2}$ |

Supp. Table II: Disequilibrium Model Constraints (Continued)

| | **No AM, No Latent PGS's** | **No AM, Latent PGS's** | **AM, No Latent PGS's** | **AM, Latent PGS's** |
|---|---|---|---|---|
| cov($Y_p$, $L[N]T_m$)<br>cov($Y_m$, $L[N]T_p$) | – | 0 | – | $V_{Y1}\mu\Gamma_1$ |
| cov($Y_p$, $[N]T_m$)<br>cov($Y_m$, $[N]T_p$) | 0 | 0 | $V_{Y1}\mu\Omega_1$ | $V_{Y1}\mu\Omega_1$ |
| cov($Y_p$, $F_m$)<br>cov($Y_m$, $F_p$) | 0 | 0 | $V_{Y1}\mu(\delta w_1 + V_{F1})$ | $V_{Y1}\mu(\delta w_1 + av_1 + V_{F1})$ |
| cov($F_p$, $\mathrm{F}_m$) | 0 | 0 | $\mu(\delta w_1 + V_{F1})^2$ | $\mu(\delta w_1 + av_1 + V_{F1})^2$ |
| cov($Y_p$, $Y_m$) | 0 | 0 | $V_{Y1}^2\mu$ | $V_{Y1}^2\mu$ |
| cov($Y_o$, $Y_*$) | $\delta\Omega_1 + fV_{Y1}$ | $\delta\Omega_1 + a\Gamma_1 + fV_Y$ | $(\delta\Omega_1 + fV_{Y1})(1 + V_Y\mu)$ | $(\delta\Omega_1 + a\Gamma_1 + fV_{Y1})(1 + V_Y\mu)$ |
| cov($Y_*$, $F_o$) | $fV_{Y1}$ | $fV_{Y1}$ | $fV_{Y1}(1 + V_{Y1}\mu)$ | $fV_{Y1}(1 + V_{Y1}\mu)$ |
| cov($Y_*$, $F_*$) | $\delta w_1 + V_{F1}$ | $\delta w_1 + av_1 + V_{F1}$ | $\delta w_1 + V_{F1}$ | $\delta w_1 + av_1 + V_{F1}$ |

Parameter constraints under disequilibrium. As in Supp. Table I, we use the $*$ subscript to denote either $p$ or $m$ but not both within a single term. We also use subscripts 1 and 2 to denote parent and offspring generations, respectively. For example, $g_1$ represents the covariance within each parental haplotype (the "cis" correlation; e.g., $cov(T_p, NT_p)$), while $g_2$ represents the covariance between parental haplotypes (the "trans" correlation; e.g., $cov(T_p, T_m)$).
